# Genetic assimilation and accommodation shape adaptation to heat stress in a splash pool copepod

**DOI:** 10.1101/2025.09.09.675189

**Authors:** Isabelle P. Neylan, Rujuta V. Vaidya, Emma L. Crable, Brant C. Faircloth, Maheshi Dassanayake, Morgan W. Kelly

**Author notes:** **Author Contributions:** IPN, MWK conceived the ideas and designed methodology; IPN, RVV, and ELC collected the data; IPN and RVV analyzed the data; IPN and MWK led the writing of the manuscript; BCF and MD contributed critically to revisions; All authors reviewed the paper and gave final approval for publication. **Competing Interest Statement:** The authors declare no competing interest.

## Abstract

Understanding how organisms respond to variable environments is becoming increasingly important in our rapidly changing world. Beyond genetic adaptation, plastic responses to the environment can alter phenotypes and fitness, ultimately driving evolution. However, the interaction between plasticity and adaptation during environmental change is complex and hard to measure in natural systems. Here, we used two populations of *Tigriopus californicus* copepods, a thermally tolerant southern population and a thermally sensitive northern population, to conduct a fully factorial split brood experiment where we exposed animals as larvae and adults to either a sublethal heat stress or control (no heat treatment) before measuring heat tolerance and gene expression patterns. We found that increased thermal tolerance across populations came at the expense of physiological plasticity and evolved through higher baseline expression of heat stress response genes across environmental contexts as well as increased gene expression plasticity in response to heat stress. In the thermally sensitive northern population, developmental exposure to heat stress led to higher adult tolerance and lower physiological plasticity underpinned by higher gene expression plasticity. Importantly, we found that the same set of genes were largely responsible for both the evolved higher tolerance in the southern population and the developmentally induced tolerance in the northern population suggesting that in this system, a shared molecular response contributes to acclimation and adaptation across both populations. These results link existing physiological plasticity with long-term evolutionary responses providing insight into how these populations will adapt and respond to future environmental change.

**SIGNIFICANCE:** Understanding plastic and evolutionary responses to dynamic environments is critical to anticipating species’ vulnerability to climate change. In this study, we compared gene expression and physiological responses to heat stress across two populations of a marine copepod that differ in thermal tolerance to investigate mechanisms of adaptation. We found evidence for plasticity-led evolution in this system, with the same set of genes contributing to long-term evolutionary changes across populations and to short-term physiological adjustments within populations. Our results suggest that populations with a reservoir of plasticity have a greater potential to evolve as the climate continues to warm, but that there may be a limit to this adaptive capacity.

## INTRODUCTION

How species adapt to changing environments is a key problem in evolutionary biology and is becoming increasingly critical to understand in our rapidly changing world (1, 2). Evolutionary responses occur via genetic adaptation, but a single genome can produce multiple phenotypes in response to the environment, meaning that phenotypic plasticity can also play an important role in determining which individuals successfully adapt and persist (3, 4). However, the interplay between plasticity and adaptation during environmental change is complex (5, 6). Plasticity may mask underlying genetic variation, slowing responses to selection (7) or plasticity may facilitate adaptation by allowing populations to persist long enough to allow for evolutionary rescue (8, 9). But most intriguingly, the evolution of plasticity itself can pave the way for subsequent adaptation by facilitating the production of genetic variation in environmentally responsive traits (10–12). To predict future evolutionary outcomes in changing environments, it is critical to understand the role plasticity plays in genetic adaptation over time as well as the underlying mechanisms that promote or suppress plasticity in a population (13).

Gene networks that underlie plastic traits are built by many regulatory and protein coding changes over long evolutionary timescales (14, 15). But once these networks exist, changes in gene regulation can create variation in the environmental responsiveness of a trait (16), which is, in turn, the raw material necessary for further evolutionary changes in plasticity (17, 18). In variable or changing environments, selection for better environmental tracking should favor the evolution of increased plasticity, a process known as genetic accommodation (5, 19). Over time, if the environment becomes stable or if plasticity becomes maladaptive, selection will favor the loss of environmental responsiveness via the process of genetic assimilation (20, 21). Traits with greater plasticity and environmental sensitivity tend to harbor greater genetic variation and thus a higher capacity for evolutionary change (22, 23). Therefore, evolutionary changes that lead to reduced plasticity, such as genetic assimilation, can increase vulnerability to changing environments by limiting genetic variation.

The magnitude of plasticity can be modulated by the timing of exposure to a given cue. Many of the most well-studied plastic traits are irreversible and triggered by cues encountered during development (24–26). But reversible acclimation throughout adulthood also plays a role in adaptive evolution, especially in response to rapid environmental shifts, and both forms of plasticity interact to shape adult phenotypes and fitness (20, 27). Developmental priming can heighten or dampen the environmental sensitivity of a trait during adulthood, an interaction ultimately shaped by changes in gene expression (28). Early exposure to a cue may result in upregulation of environmentally responsive genes during development that are maintained at a higher baseline into adulthood, resulting in carryover effects of early exposure and the loss of adult plasticity. On the other hand, early exposure may alter transcriptional machinery to produce higher plasticity in adult gene expression and increase the environmental sensitivity of a trait (29). Over time, selection can act on these same gene regulatory modifications to drive the process of genetic accommodation or assimilation (30).

Evaluating the contribution of genetic accommodation and assimilation to adaptation is notoriously difficult in natural systems where it is only possible to measure end points of evolution (31). One approach is to use a space-for-time substitution by measuring the means and plasticities of environmentally responsive traits across populations that are locally adapted to a known environmental gradient (32). In these cases, phenotypic measurements can be combined with transcriptomic data to investigate whether loci that are environmentally responsive within populations are also disproportionately involved in population divergence (33, 34). In environmental change studies, this involves comparing a more “tolerant” population to a more “sensitive” population and exposing each to control and stress conditions to compare phenotypic and molecular responses (29, 35). If stress tolerance has evolved through genetic accommodation, we expect the tolerant population to show greater gene expression plasticity in environmentally responsive genes after exposure to the stressor than the sensitive population. If stress tolerance has evolved through genetic assimilation, we expect the tolerant population to have a higher baseline expression of beneficial genes in control conditions and to have less gene expression plasticity when exposed to the stressor (36).

*Tigriopus californicus* copepods are a tractable, well-established system in molecular ecology and physiology and well-suited for environmental change studies. *T. californicus* occupies high intertidal splash pools along the West coast of North America from Baja California to Alaska, where they experience substantial latitudinal, seasonal, and diurnal variation in temperature (37). These copepods have short generation times (about three weeks) and limited gene flow, resulting in genetically diverged and locally adapted populations (38). Importantly, *T. californicus* populations vary in their ability to thermally acclimate as adults after exposure to a sublethal heat shock (39, 40) as well as how exposure to stressful temperatures during larval development impacts subsequent heat tolerance in adulthood (41, 42). Here, we used two populations of *T. californicus* from along the coast of California, USA — a thermally sensitive northern population and a thermally tolerant southern population (see Materials & Methods). We measured phenotypic (thermal tolerance) and transcriptomic (differential gene expression) responses to temperature stress experienced during early-life and adulthood across the two populations using a fully factorial design. Our goals were to address: 1) how exposure to heat stress across life stages impacts thermal tolerance and plasticity, 2) whether the heat tolerant southern population gained an increased tolerance through evolution of plasticity via increased trait mean and loss of plasticity (genetic assimilation) or through the evolution of increased acclimation potential (genetic accommodation) by comparing patterns of differential gene expression and to determine, 3) which genes are involved in phenotypic plasticity and evaluate how the environmental responsiveness of these genes was modified in the tolerant southern population.

## RESULTS

### Thermal tolerance assays reveal higher tolerance but lower physiological plasticity in southern population

We performed thermal tolerance assays on adult copepods that were originally collected from two wild populations (northern and southern California) but cultivated in constant lab conditions for > 15 generations and exposed to one of four treatments: 1) control (no sublethal heat shocks), 2) adult shock, 3) larval shock, and 4) larval + adult shock (Fig. 1A). To calculate upper lethal limits, we constructed a logistic regression from survival data measured across a range of temperatures to estimate a LT50 value, or the temperature (°C) at which 50% of animals died within each treatment (26, 37). As expected, the southern population had a higher tolerance (37.6 ± 0.04° C for southern controls compared to 35.8± 0.04° C for northern controls [mean LT50 ± SE]; Fig. 1B). We also found that the northern population had a stronger adult acclimation effect than the southern population (an increase of 0.6° C versus 0.3° C) although both populations demonstrated increases in tolerance after a sublethal adult shock (Fig. 1B; solid lines). We did not find an effect of larval shock in the southern population, but we observed a significant effect of larval heat shock in the northern population (Fig. 1B; dashed lines). Specifically, a larval shock in the northern population increased tolerance almost as much as an adult shock alone, but there was no additional benefit to being shocked as a larva and again as an adult.

**Figure 1.**
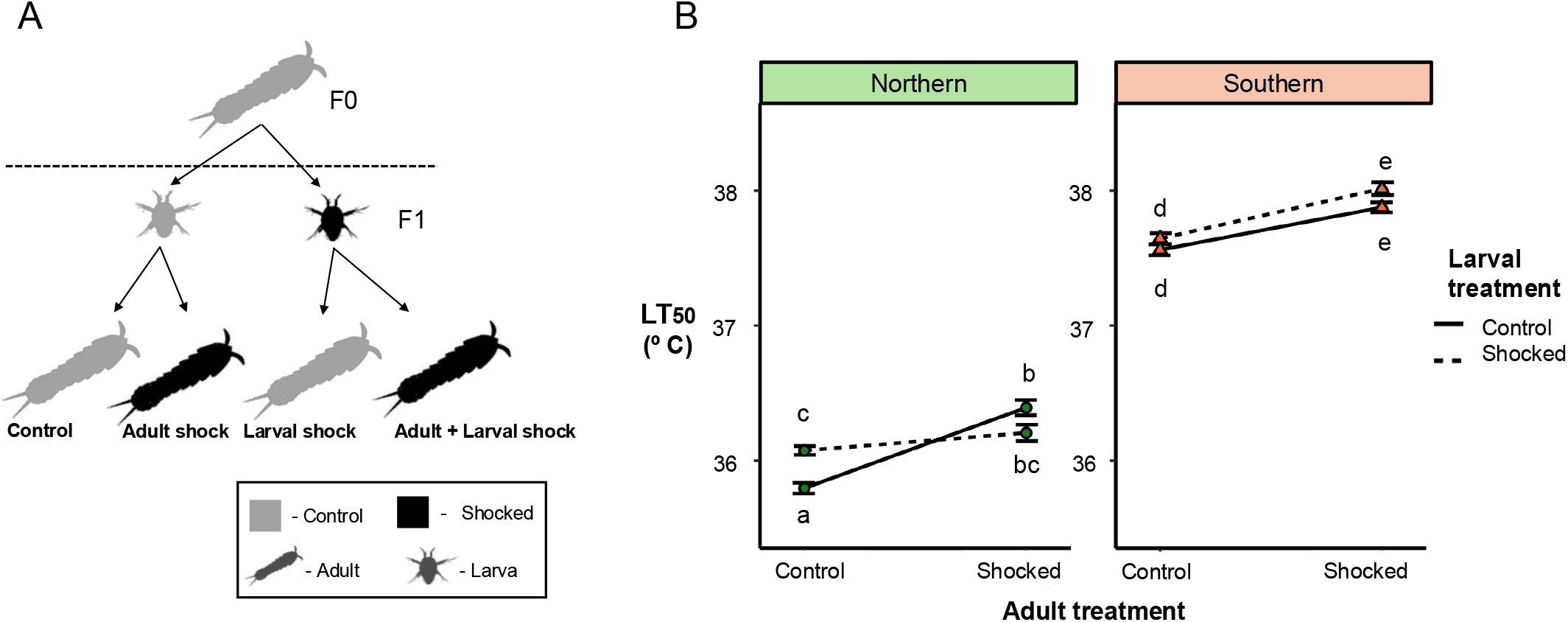
A) A schematic illustrating the experimental design for northern and southern *Tigriopus californicus* populations. Larvae were collected from parents maintained in control conditions and half were exposed to a sub-lethal but stressful heat shock (shocked, black) for one hour while the other half were maintained in control conditions (control, gray). These larvae were allowed to grow into adulthood where half of the adult animals were exposed to a sub-lethal heat stress (shocked) for one hour while the other half remained in control conditions (control) for a fully factorial design. B) The results of the LT50 heat tolerance assays for both the northern population (left panel) and southern population (right panel). Values along the y-axis represent the temperature (ºC) at which 50% of the animals in each treatment group died. Adult treatment is indicated across the x-axis (control or shocked) and larval treatment is indicated by the line type (solid indicates control, dashed indicates shocked). Different letters represent significantly different values between treatment groups (non-overlapping 95% confidence intervals around the LT_50_ estimate) and error bars represent standard error.

### Gene expression comparisons suggest higher tolerance in the southern population evolved via accommodation and assimilation

We assessed gene expression responses in the northern and southern populations by comparing adults exposed to heat shock with their respective control groups. Our goal was to identify gene expression changes associated with heat stress acclimation observed in both populations and to explore the mechanisms driving higher tolerance in the southern population. To obtain a candidate list of heat responsive genes, we conducted four differential gene expression analyses: 1) between northern control and northern adult shock treatments, 2) between northern control and northern larval + adult shock treatments, 3) between southern control and southern adult shock treatments, and 4) between southern control and southern larval + adult treatments. Through these analyses, we found 485 genes significantly upregulated in response to heat stress across our two populations (*p*-adjusted < 0.05; Table S1; Supplemental Methods). We then compared TPM (transcripts per million) values for each of the 485 genes and categorized each gene depending on: 1) whether it had higher baseline expression in the absence of heat stress in southern controls than in northern controls, which would provide evidence for genetic assimilation, and 2) whether the increase in expression after heat shock was higher in the southern than the northern population (i.e., whether there was a greater plastic response in gene expression in the southern population), which we interpreted as evidence for genetic accommodation (Table S2).

We found that 163 genes supported a pattern of genetic assimilation, meaning that they had a higher baseline expression in southern control than northern control animals and did not show increased expression after adult shock in the southern population (Fig. 2). Using a gene ontology enrichment analysis, we determined that these genes were associated with peptidases (17 genes *p* ≤ 0.06), oxidoreductases (21 genes, *p* < 0.001), and chitin binding (9 genes, *p* < 0.001; Fig. 2C; Table S3). We found 85 genes that supported a pattern of accommodation, meaning that they were not constitutively upregulated in southern controls, but had a greater increase in expression after adult heat stress in the southern as compared to the northern population. These genes were enriched for heat shock proteins and protein folding processes (10 genes, *p* < 0.001; Fig. 2C; Table S4). 148 genes had higher baseline expression in southern than northern controls but were also more plastic in the southern population after heat shock, meaning that these genes retained their plasticity and better support a pattern of accommodation rather than assimilation. These genes were significantly enriched for heat shock proteins and protein folding processes (13 genes, *p* < 0.05) as well as peptidases and proteolysis (26 genes, *p* ≤ 0.07; Fig. 2C; Table S5). Finally, 89 genes did not support a pattern of either genetic assimilation or genetic accommodation because they showed lower baseline expression and lower plasticity in the southern population. We found this category to be enriched for genes related to antioxidants and oxidoreductases (12 genes, *p* β 0.05), and chitin binding (6 genes, *p* < 0.001; Fig. 2C; Table S6).

**Figure 2.**
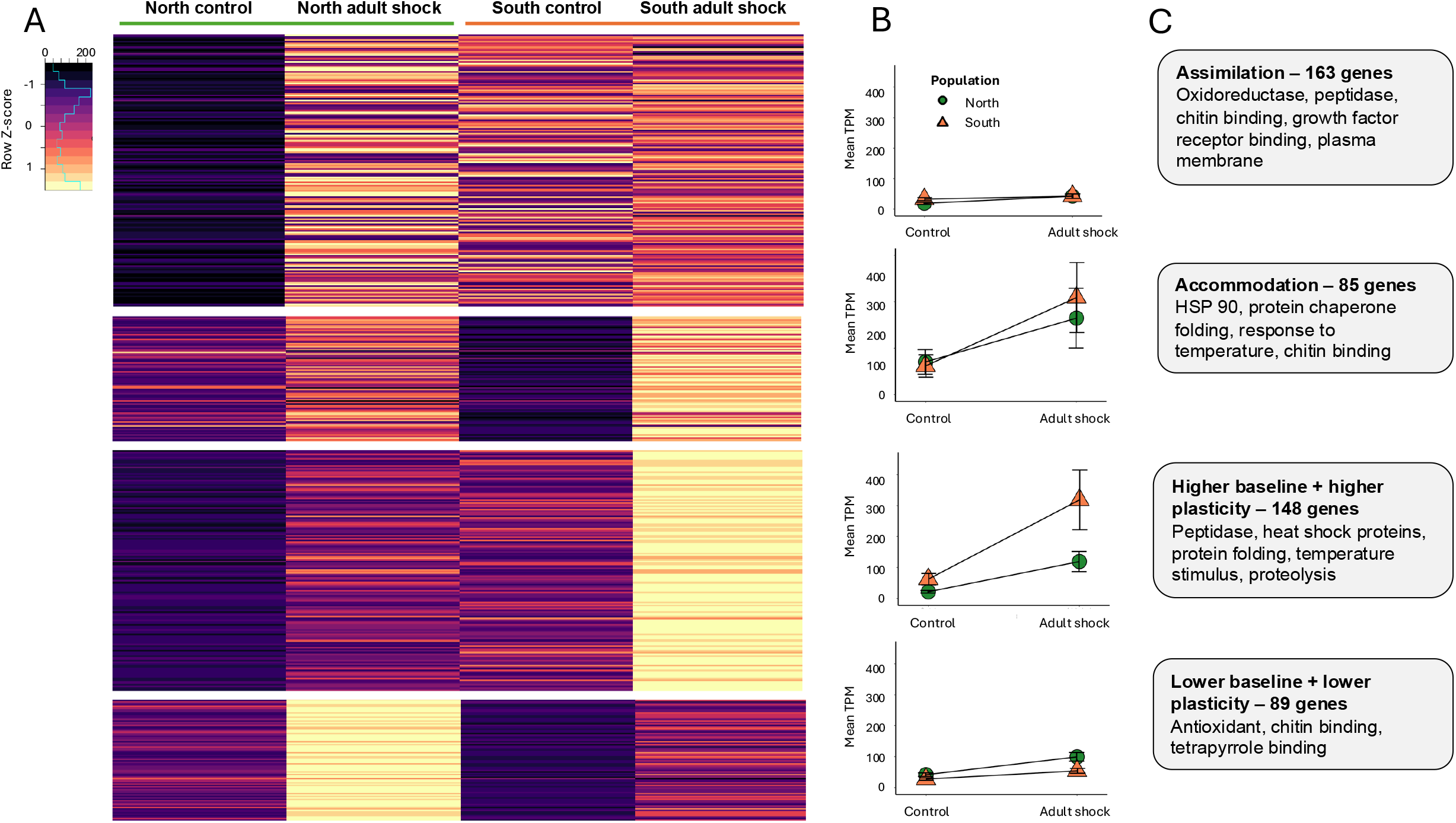
Comparison of the differentially expressed genes in response to adult heat stress in the northern and southern *Tigriopus californicus* populations. A) Expression heat map of the 485 genes that were significantly up regulated in response to adult heat stress across northern and southern copepods. The mean TPM (transcripts per million) value was used for each population and treatment combination (calculated using three biological replicates) to calculate a relative expression value (Z-score). The categories were determined by comparing 1) the expression level of each gene in the southern control to the northern control (i.e., whether the southern population had higher baseline expression in controls, evidence for assimilation), and 2) the difference in expression between southern control and adult shock and northern control and adult shock treatments (i.e., the difference in slope, indicating higher plasticity or accommodation). The numbers under each category in parentheses indicate the number of genes assigned to the category. B) Reaction norm plots illustrating the mean expression level (measured in TPM) within each category and across the four populations and treatment combinations. Control and adult shock along the x-axis indicate treatment and color and shape indicate population (green circles indicate northern population, orange triangles indicate southern population). Error bars indicate standard error. C) A summary of the category, the number of genes within the category, and the gene ontology enrichment analysis indicating terms were significantly enriched within that category.

### Developmental priming in the northern population leads to increased transcriptional plasticity in adulthood

We followed a similar framework to explore the mechanism behind the larval priming effect we found in the northern population. Using our differential gene expression analyses in the northern population between control and adult shock as well as control and larval + adult shock animals, we identified 419 heat responsive genes and compared TPM values across the four treatment combinations (control, adult shock, larval shock, and larval + adult shock) within the northern population. We categorized each gene based on: 1) whether expression was higher in adult animals that experienced heat shock as larvae compared to control animals when measured in control conditions, evidence for developmental priming leading to maintenance of higher baseline expression, and 2) whether there was increased plasticity between control and adult shock treatments if animals had received a larval shock (Table S7). We ran the same analysis for the southern population using a list of southern heat responsive genes (186 genes) and confirmed that the lack of larval shock effect in thermal tolerance was accompanied by a lack of strong gene expression patterns (see Supplemental Methods; Fig. S1; Table S16-18).

In our northern population, we found 59 genes that demonstrated a pattern of higher baseline expression without an increase in plasticity, but we did not find significant gene enrichment within this category (Fig. 3A, B, C). 204 genes showed higher plasticity but not higher baseline expression in animals receiving a larval shock, and this category was significantly enriched for genes related to the heat shock protein response and protein folding processes (14 genes, *p* < 0.001; Fig. 3C; Table S8). 138 genes demonstrated higher baseline expression and higher plasticity and were enriched for categories related to peptidases and proteolysis (22 genes, *p* < 0.001) as well as oxidoreductase and antioxidants (19 genes, *p* β 0.05; Fig. 3C; Table S9). Finally, the remaining 18 genes showed similar responses to adult heat shock between larval treatments, and we were unable to identify any enriched genes in this category.

**Figure 3.**
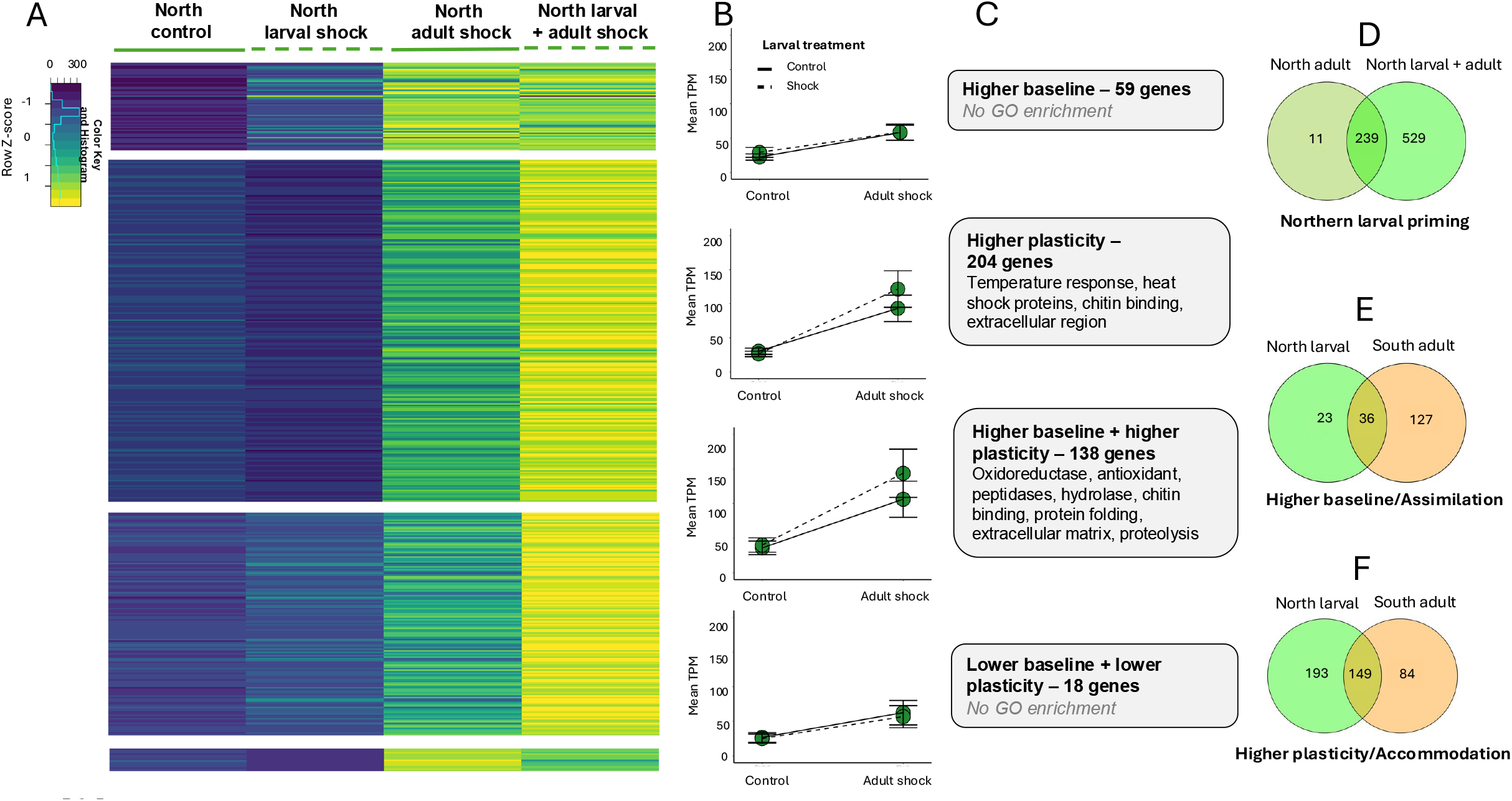
Comparison of the differentially expressed genes in the northern *Tigriopus californicus* population across four treatments (control, larval shock, adult shock, and larval + adult shock). A) Expression heat map of the 419 genes that are significantly up regulated in copepods exposed to adult heat shock as compared to controls. The mean TPM (transcripts per million) value was used for each population and treatment combination (calculated using three biological replicates) to calculate a relative expression value (Z-score). The categories were determined by 1) comparing the expression level of each gene in the larval shock treatment to the control treatment (i.e., if the larval shock treatment had a higher baseline expression) and 2) the difference in expression between control and adult shock and larval shock and larval + adult shock treatments (i.e., the difference in slope, indicating higher plasticity). B) Reaction norm plots illustrating the mean expression level (measured in TPM) within each category and across the four treatment combinations. The x-axis indicates adult treatment, and the line type indicates larval treatment (solid line indicates control, dashed line indicates larval shock). Error bars indicate standard error. C) A summary of the category, the number of genes within the category, and the gene ontology enrichment analysis indicating terms were significantly enriched within that category. D) Venn diagram displaying the significantly upregulated genes in copepods from the northern population that experienced adult shock alone or an adult + larval shock as compared to controls. E) Venn diagram displaying the genes that were considered assimilated in the southern population (Fig. 2) and genes with higher baseline expression in northern larval priming. F) Venn diagram displaying the genes that were considered accommodated or those with higher plasticity + higher baseline expression in the southern population (see Fig. 2) and genes with higher plasticity (with and without higher baseline expression) in northern larval priming.

We also wanted to better understand how the larval and adult shock treatments were interacting to affect upper lethal limits in the northern population. Using the results from our differential gene expression analyses between control and adult shock as well as control and larval + adult shock treatments, we determined which upregulated genes were unique or overlapping across our treatment comparisons (Fig. 3D). We found that while most genes upregulated in the adult shock treatment overlapped with those upregulated in the larval + adult shock treatment (96%), there were many additional genes unique to the larval + adult shock category with over three times as many significantly upregulated genes overall. Gene ontology enrichment revealed that the genes shared across both treatments were related to heat shock proteins and protein folding processes (17 genes, *p* < 0.05) while the genes unique to the larval + adult shock treatment involved additional stress responses such as genes related to peptidases and proteolysis (42 genes, *p* < 0.04) as well as antioxidants and oxidoreductase (35 genes, *p* < 0.001; Table S10-12).

### Genes responsible for evolved tolerance in the southern population largely overlap with those responsible for plasticity in the northern population

Finally, we wanted to explore whether the genes responsible for increased thermal tolerance due to evolved differences in the southern population were shared or distinct from those conferring increased tolerance due to larval priming in the northern population. Of the genes associated with constitutive responses lacking plasticity — assimilation in the southern population (162) and higher baseline expression in the northern population (59) — the northern population shared many of the individual genes with the southern population (61%) while the southern population expressed a much higher proportion of unique genes (78%) (Fig. 3E). These uniquely expressed southern genes were enriched for growth factors and receptors, chitin binding, metabolism, autophosphorylation, and cell population proliferation (Table S13). For genes associated with evolved genetic accommodation in the South (169) and higher plasticity due to developmental priming in the northern population (342), we found that the majority of genes in the southern population overlapped with those in the northern priming response (88%) and these common genes were enriched for heat shock proteins (18 genes, *p* < 0.05) as well as peptidases and proteolysis (24 genes, *p* < 0.01; Fig. 3F). Conversely, 66% of these genes were uniquely differentially expressed in the northern population and were enriched for multiple peptidases and hydrolases (22 genes, *p* < 0.06) as well antioxidants and oxidoreductase (27 genes, *p* ≤ 0.05; Table S14 and S15). Importantly, we found that the individual genes associated with assimilation and accommodation in the southern population were significantly enriched for the northern plastic response (*p* < 0.001; Fisher’s Exact test) indicating a larger overlap than we would expect by chance and suggesting that the same genes are largely responsible for evolved responses in the south and the developmentally primed heat shock responses in the north.

## DISCUSSION

We were interested in understanding the mechanistic overlap between plastic responses to heat stress within populations and evolved differences in heat tolerance between populations of *Tigriopus californicus*. If evolution of increased heat tolerance occurred through accommodation or assimilation of plastic responses, we expected the evolved responses to heat stress in the thermally tolerant southern population to resemble the plastic responses to heat stress in the thermally sensitive northern population. Consistent with these expectations, we observed similarities between evolved and plastic responses, but our inference of evolution through genetic accommodation versus assimilation depended on whether we considered the response to heat stress at a physiological or molecular level. At the physiological level, our results indicated that increased thermal tolerance in the southern population evolved through an increase in constitutive mean tolerance across environmental contexts and diminished plasticity of heat tolerance, a pattern consistent with genetic assimilation that is widespread across metazoans (40, 43, 44). Developmental priming caused the northern population to become more like the southern population as northern copepods that experienced a larval shock had higher baseline thermal tolerance and lower adult heat hardening capacity (Fig. 1B).

At the molecular level, we found evidence that the increase in mean heat tolerance in the southern population was produced by genetic assimilation and accommodation because we found transcripts showing higher baseline regulation but lower plasticity in response to heat stress (genetic assimilation) as well as transcripts showing greater plasticity in response to heat stress after an adult shock (genetic accommodation; Fig. 2). However, the most compelling evidence for genetic accommodation came from the similarities in gene expression patterns between evolved responses in the southern population and developmental plasticity in the northern population. Of the 233 transcripts with greater upregulation after heat shock in the south compared to the north, 149 also showed greater heat shock response after larval priming in the northern population, a highly significant enrichment of the *evolved* accommodation response in the southern population for genes involved in the *developmentally plastic* responses to heat shock in the northern population. Notably, gene expression patterns differed across functional categories in the northern and southern populations, with peptidases showing signatures of higher baseline expression and assimilation while heat shock proteins showed signatures of higher plasticity and accommodation.

During heat stress, peptidases can break up protein aggregations caused by misfolded or denatured proteins and help improve cellular function during stress (45, 46). Peptidases can be plastically induced in response to a stressor but can also be produced ahead of time and stored in lytic vacuoles within cells until needed (47). Consistent production of peptidases through higher baseline expression would alleviate the need for acclimation in this subset of genes and ultimately assimilation that buffers against environmental perturbation to maintain homeostasis (11). A similar pattern of transcriptional front-loading has been found between resilient and sensitive populations in their response to heat stress in several other marine species (36, 48–50).

Genes related to heat shock proteins (HSPs) were consistently associated with accommodation and higher plasticity categories in both the northern and southern responses. HSPs are conserved across the tree of life and are known to be highly inducible in response to heat stress (51, 52). However, HSPs have also been found to be a part of a front-loaded response with tolerant populations displaying higher constitutive HSP levels at baseline (36, 53). In our southern population, we found some evidence for higher baseline expression of HSPs, but this higher baseline was always accompanied by upregulation in response to heat stress, indicating that plasticity is maintained. One potential explanation for this retention of plasticity is that these genes have not had enough time or selection pressure to canalize (54). The alternative explanation is that the southern population experienced enough variability in their evolutionary history to maintain sensitivity. The preservation of plasticity may also be due to the nature of the proteins themselves. Retaining HSPs when not needed can be a maladaptive response detrimental to cellular function and thus plasticity may be maintained regardless of the selection regime (51).

Another overarching pattern across our results was a saturation of response with increased exposure to heat stress. We found decreased acclimation plasticity in the thermally tolerant southern population as compared to the northern population in response to adult shock, supporting a potential plasticity-tolerance trade-off (40, 43, 44). In the northern population, we found that the level of adult plasticity was modulated by an individual’s early-life experience, with no additional benefit of being shocked as larvae and as adults. This saturation may arise because the increase in physiological plasticity is driven by an increase in gene expression plasticity which is sensitive to thermal stress experienced during early-life and in adulthood. While we found a substantial set of differentially expressed genes that were upregulated in response to adult shock when an individual had also received a larval shock (529; Fig. 3D), the transcriptional plastic response of an adult shock, alone, was sufficient to confer similar benefits in tolerance. This result suggests that the 239 shared genes associated with HSPs and other plastically induced responses were sufficient to confer increased tolerance and may have already reached maximum transcriptional plasticity such that two shocks (larval + adult) provided no additional increase in transcriptional plasticity above and beyond that generated by a single shock. Theoretical and modeling studies suggest that populations that have evolved with a more variable environmental regime and display higher levels of plasticity are better able to persist long enough for evolutionary rescue in the face of environmental change (55–57). The retention of plasticity and lack of canalization in multiple genes in the northern and southern populations suggests that there may be room for further adjustment as temperatures increase or continue to vary, but there may be a hard limit to the amount of heat tolerance these plastic responses can confer due to this pattern of saturation.

Overall, the mechanisms underlying the adaptive responses in the southern population were largely shared with the larval acclimation effect in the northern population, linking current physiological plasticity with long-term evolutionary responses and providing evidence for plasticity-led evolution. Our empirical data support the theory that physiological responses to the environment can respond rapidly to selection because environmental stressors are heterogenous and frequently vary over generational time (58) and that selection should act on genes with environmentally responsive expression because these genes underlie phenotypes relevant to different fitness optima across environments (22, 23). In this system, adaptation to environmental change appears to be occurring through modification of shared regulatory networks as evidenced by the differences in expression of environmentally responsive transcripts between the northern and southern population underpinned by shared genetic pathways (27, 34). The ability to adjust the environmental responsiveness of gene expression and ultimately alter phenotype in response to environmental variation appears to be a driver of adaptive evolution in this system. Our results suggest that the northern population with its larger reservoir of plasticity should have a greater potential to evolve as the climate continues to warm, but that there is a limit to this adaptive capacity which the southern population appears to be approaching. Further understanding about how trait innovation and stress tolerance evolves across a diversity of taxa will improve our ability to predict species and population persistence and evolutionary potential into the future.

## MATERIALS & METHODS

### Animal collection and experimental design

During the Summer of 2022, we collected *Tigriopus californicus* copepods from two field sites in California USA: Salt Point State Park near Jenner, CA (Northern population; 38° 33’ 58.34” N, −123° 19’ 55.32” W) and Sunset Cliffs in San Diego, CA (Southern population; 32° 43’ 56.83” N, −117° 15’ 24.04” W) and used these to establish lab cultures following procedures described in the Supplemental Methods and elsewhere (37, 40, 59). Using these animals, we ran a fully crossed factorial, split-brood experiment with animals experiencing either a sublethal heat shock (34° C for 1hr) or maintained at the control temperature (19° C) at both an early-life larval stage (larval exposure) and/or as an adult 24-48 hours prior to thermal tolerance experiments (adult exposure) (Fig. 1A). We implemented this design across both populations and measured adult thermal tolerance as well as transcriptomic gene expression responses.

### Thermal tolerance assays

Within 24-48 hours after the adult heat shock or control treatment, copepods from all four treatment combinations were subjected to our thermal tolerance assay designed to measure lethal temperatures and calculate an LT50 value or the temperature at which 50% of animals die (37, 60). We exposed at least three replicate tubes containing six animals to each target temperature for a total of 348-534 animals tested per treatment by population combination (3,852 animals total; See Supplemental Methods for additional details). The assay consisted of a 1hr acclimation period at 30° C before ramping up to the target temperature for an additional hour. Animals were allowed to recover for 24hrs before we counted the number of survivors in each tube. We calculated the LT50 temperatures and their associated error terms for each treatment and population combination by fitting logistic regression models to the binomial survival data using the *drc* package in R (61).

### RNA extraction and differential gene expression analysis

We flash froze a separate subset of adult copepods using liquid nitrogen immediately after experiencing the sublethal adult shock or control treatment. Using pooled samples of 50-60 copepods per replicate, we extracted total RNA using steps 1-7 of the TRIzol (Invitrogen) protocol followed by steps 4-11 of the Qiagen RNAeasy Plus kit (Qiagen) protocol. We sent samples to Novogene Corporation Inc. in Sacramento, California to create a total of 24 libraries (three per population and treatment combination) that were sequenced on NovaSeq X Plus with 150-bp paired-end reads. We mapped the resulting sequence reads to the *T. californicus* reference genome (62) using *STAR* RNA-seq aligner (version 2.6.0a) (63) and computed transcripts per million (TPM) values using the *RSEM* package (64). We conducted differential gene expression analyses using *DESeq2* (v 1.24.0) (65) and gene ontology enrichment analyses using *GO_MWU* (66) in R version 4.0.3 (67).

## Supporting information

Supplemental Results

