## Supplemental Methods for "Genetic assimilation and accommodation shape adaptation to heat stress in a splash pool copepod"

*Animal collection and maintenance*

We collected *Tigriopus californicus* animals from the field and used these to establish lab cultures from two sites in California USA: Salt Point State Park near Jenner, CA (Northern population; 38° 33' 58.34" N, -123° 19' 55.32" W) and Sunset Cliffs in San Diego, CA (Southern population; 32° 43' 56.83" N, -117° 15' 24.04" W) in the Summer of 2022. For additional details on field collection and culturing see (1–3). We chose these two populations because they have well-documented differences in mean thermal tolerance and thermal plasticity (1, 3). We maintained our lab cultures in an incubator (Percival 136VL) at 19° C under a 12hr. light/12hr. dark cycle and using artificial seawater (Instant Ocean®) at a salinity of 35ppt for at least 15 generations before the start of these experiments. All animals were fed ground up *spirulina* fish flakes *ad libitum*. To reach adequate sample sizes for our thermal tolerance assays, we repeated our experimental procedure five times to create five temporal replicates spaced about a week a part with collection of the first six larval stages (nauplii) running from September to October of 2023. We repeated the same procedure one more time in May of 2024 using 50 gravid females (F0) to create the F1 animals used in our transcriptomic experiments.

*F1 larval exposure*

We created the F1 generation by collecting 50 gravid females from our stock containers (F0) and placing each individual female into one well of a six-well plate and allowing her to hatch multiple broods of nauplius larvae over the course of two weeks. Using a dissecting scope and glass pipette, we carefully transferred half of each brood into four 200mL, lidded plastic cups filled with 150ml of seawater (representing each of our four larval-adult treatment combinations) to ensure that every mother contributed to the treatments equally (split-brood design). Each cup housed 200-350 nauplii (except for replicate 1 that only had about 150) and mothers typically produced between 0 and 50 nauplii.

After the nauplii were collected, we placed two of the four cups into a water bath (VWR WBE20 General Purpose Water Bath) set at 34° C for 1 hour to serve as our sublethal heat shock while we maintained the other two as room temperature controls. This sublethal shock served as our larval exposure. After the hour, we removed the cups from the bath and maintained all treatment cups in the same conditions as our lab cultures (in an incubator at 19° C, 12hr. light/dark cycle, 35 ppt) until the copepods reached adulthood (past final molt stage).

*F1 adult exposure*

Once animals had reached adulthood (about two weeks later), we transferred six animals into 0.6mL PCR tubes filled with 200µL of seawater at 35ppt to create as many replicates as we could for the thermal tolerance assay (n = 6-27 tubes per temporal replicate per population and larval-adult treatment combination; A small number of tubes ended up having more or less than six animals (4-9 animals)). We used males and females and included mate-guarding pairs but excluded gravid females and any copepods that had not matured to adulthood at the time of the assay.

We placed half of the tubes into the water bath for 1 hour at a temperature of 34°C while the other half remained at room temperature to create our four treatment combinations spanning our larval and adult treatments (Control-Control (control), Shock-Control (larval shock), Control-Shock (adult shock), and Shock-Shock (larval + adult shock)). While there was some background mortality during the maturation of the copepods, these rates were low and did not significantly differ across our treatments confirming that our larval shock was sublethal (4). We did not see any mortality during the period between the adult shock and the thermal tolerance assay as found in previous studies (1, 3, 5).

*Thermal tolerance assays*

Within 24-48 hours after the adult heat shock, we subjected tubes from all four treatment combinations to our thermal tolerance assay designed to measure lethal temperatures and calculate an LT50 value or the temperature at which 50% of animals die (1, 6). We exposed three replicate tubes with six animals each (18 total) to each target temperature. We began our assays at the estimated LT50 found in previous studies from our lab that included these two populations (North control – 35.6; North heat shocked – 38.0; South control – 37.2; South heat shocked – 37.8) (1) and then progressing by 0.2° C up and down from that point until we found temperatures with 100% mortality and 0% mortality (13-16 temperatures tested per population and treatment combination). We ran additional replicates for temperatures that produced around 50% mortality to increase accuracy for these critical temperatures (n = 3-16 tubes or 18-96 animals per temperature across populations and treatments). The assay consisted of a 1hr acclimation period at 30° C before ramping up to the target temperature for an additional hour within a thermocycler to precisely control temperatures (Techne 3PRIMEX/02, Bibby Scientific LTD) (1, 3). Animals were allowed to recover for 24hrs before we counted the number of survivors in each tube.

*Thermal tolerance statistical analyses*

We performed all analyses in R version 4.4.0 (7). We calculated the LT50 for each treatment and population combination using the *drc* package (Ritz et al., 2015). For all treatments across both populations and across all five temporal replicates, we fit a logistic regression model (LL.2) that allowed us to estimate an LT50 value as well as the 95% confidence interval and standard error associated with this estimate. We corrected for simultaneous inferences using the *glht* (for standard error) and *confit* (for confidence intervals) functions from the *multcomp* package (8). We considered treatments to have significantly different LT50 values ($p < 0.05$) when their 95% confidence intervals did not overlap (6, 9)

*Tissue collection and RNA extraction*

We used a different subset of copepods for our RNA extractions but followed the same protocols for F1 nauplii collection and early-life larval exposure as the thermal tolerance assay animals (see above). We maintained the animals at control conditions in the incubator until maturation before we began gene expression experiments on the adult animals. We placed copepods from the larval control and larval shock treatment into 50mL falcon tubes filled with seawater and exposed these to either: 1) control, animals that did not experience any adult heat shock, or 2) shock, animals were exposed to a sublethal adult shock at 34° C for 1hr and immediately frozen. We subdivided each 50mL falcon tube into four replicate 1.5mL Eppendorf tubes with 50-60 animals in each (n = 32 tubes total, four replicates for each population and treatment combination). We flash froze all animals using liquid nitrogen and stored them in 1.5mL falcon tubes in a -80° C freezer until extraction.

*RNA extraction and sequencing*

We extracted total RNA from flash frozen copepods using a combination of TRIzol (Invitrogen; catalog no. 15596026) and the Qiagen RNAeasy Plus kit (Qiagen, catalog no. 74134). We homogenized copepods using a TissueRuptor II (Qiagen, catalog no. 9002755) before we followed steps 1-7 of the TRIzol protocol, followed by steps 4-11 of RNAeasy Plus kit protocol (10). We extracted total RNA from 24 samples and sent extracts to Novogene Corporation Inc. (Sacramento, California), where RNA quality was confirmed using a 2100 Agilent Bioanalyzer on a Eukaryote Total RNA Nano chip followed by the preparation of non-directional RNASeq libraries using poly-A tail selection. The resulting libraries were sequenced on NovaSeq X Plus, with 150-bp paired-end reads.

Sequencing quality was checked using the program FASTQC (11). We removed adapter sequences and low-quality reads using Trimmomatic (12). We then mapped the reads to the *T. californicus* reference genome (13) using STAR RNA-seq aligner (version 2.6.0a) (14). Reads were mapped to gene features with the options (--quantMode GeneCounts --outFilterScoreMinOverLread 0.50 --outFilterMatchNminOverLread 0.50) to adjust for poly-A tail contamination (15), which generates a count matrix using ReadsPerGene.out.tab output. We generated transcripts per million (TPM) values used for visualizing and analyzing our RNASeq dataset using the RSEM package (16).

*Differential gene expression analysis*

We removed genes that were expressed at a low level, which we defined as genes that did not have at least 10 counts in 25% of samples (17) from our dataset. This retained 16,821 genes that we used in subsequent analyses. We conducted differential gene expression analyses using DESeq2 (v 1.24.0) (18). For each pairwise comparison we obtained a list of differentially expressed genes (DEGs) and calculated false discovery rates (FDR) using the Benjamini–Hochberg method (19). We considered genes with an adjusted $p$-value cutoff of $< 0.05$ and a log2foldchange of $> 1$ (for upregulated genes) or $< -1$ (for downregulated genes) to be significantly differentially expressed (see Table S1 for all counts of up- and down-regulated genes per pairwise comparison). We performed gene ontology enrichment using GO_MWU (20), using the Fisher's Exact Test ($p < 0.05$) and the scripts available at: https://github.com/z0on/GO_MWU/blob/master/GO_MWU.R. We used TPM values to create heatmaps using the *heatmap.2* function in the *gplot* package (11) and the reaction norm graphs using the *ggplot2* package (21). We constructed Venn diagrams using the DEG counts determined through our various pairwise comparisons and visualized these using the *ggvenn* function in the *ggplot2* package.

*Analysis of North versus South evolved gene expression responses*

To understand the potential differences between our more thermally tolerant southern population and more thermally sensitive northern population, we began by conducting a differential gene expression analyses using the following pairwise comparisons: control copepods compared to those that had been shocked as adults (adult shock) in the northern population, control copepods compared with those shocked as adults and as larvae (larval + adult shock) in the northern population, control copepods compared to adult shock in the southern population, and control copepods compared with larval + adult shock copepods in the southern population. These analyses resulted in a combined list of 485 significantly up-regulated differentially expressed genes (DEGs) found across the four pairwise comparisons (Table S2). Including both adult shock treatments was important because it allowed us to examine the full suite of genes potentially involved in the heat stress response across contexts.

To place these genes within each of our hypothetical categories (assimilation, accommodation, higher baseline + higher plasticity, lower baseline + lower plasticity), we created two true-false binaries and compared the averaged TPM within a treatment (i.e., across the three replicates). The first asked whether the baseline expression level of these DEGs in southern controls was greater than that in northern controls, which was considered potential evidence of assimilation. The second asked whether the difference between control and adult shock in the southern population was greater than the difference in the northern population, evidence of increased plasticity or accommodation. Genes that showed evidence for being true for both were considered to support a “higher base line + higher plasticity” hypothesis. Genes that were true for the first and false for the second were considered evidence for “assimilation”. Genes that were false and true were considered evidence for “accommodation”. Finally, genes that were false for both did not fit into any of those categories (“lower baseline + lower plasticity”).

*Analysis of Northern developmental priming through gene expression*

We used the results of the differential expression analysis between control vs. adult shock animals and control vs. larval + adult shock animals in the northern population to create a list of 419 northern heat responsive genes to conduct our analysis of the larval priming mechanism in the northern population. We constructed two true-false binaries and compared TPM values, but this time across the four treatments in the northern population – control, adult shock, larval shock, and adult + larval shock. The first binary asked whether the expression level of a given gene in the larval shock treatment was higher than in controls (no adult shock for either), evidence of higher baseline expression. The second asked whether the difference between control and adult shock was higher in copepods that had a larval shock (larval shock vs. larval + adult shock) than those that lacked one (control vs. adult shock), evidence for increased plasticity after an adult shock. This allowed us to assign each gene to one of our four categories (higher baseline only, higher plasticity only, higher baseline + higher plasticity, and lower baseline + lower plasticity). We also used this list of DEGs to better understand the mechanistic differences induced by larval priming by constructing the Venn diagram in Fig. 3D and comparing common and unique genes across the northern treatments.

*Analysis of Southern developmental priming through gene expression*

Finally, despite little evidence for larval priming effects in the southern population (i.e., no difference in thermal tolerance (Fig. 1B), zero DEGs between southern control copepods and southern larval shock copepods, and little difference in the number of DEGs between the pairwise comparisons of control vs. adult shock (227) and control vs. larval + adult shock (258; Table S1)), we wanted to ensure our conclusions were complete so we performed a similar developmental priming analysis as we did for the northern population (described above). Using the results from the differential expression analyses comparing control vs. adult and control vs. larval + adult shock treatments in the southern population, we created a list of 186 significantly upregulated genes responsive to heat shock (Table S16). We used the same binary framework as with the northern population to determine which genes belonged in each of our four categories (higher baseline only, higher plasticity only, higher baseline + higher plasticity, and lower baseline + lower plasticity; Fig. S1; Table S21-24). Ultimately, genes were evenly distributed across our four categories without a clear signature of higher plasticity or higher baseline expression.
