## Supplemental Results for "Genetic assimilation and accommodation shape adaptation to heat stress in a splash pool copepod"

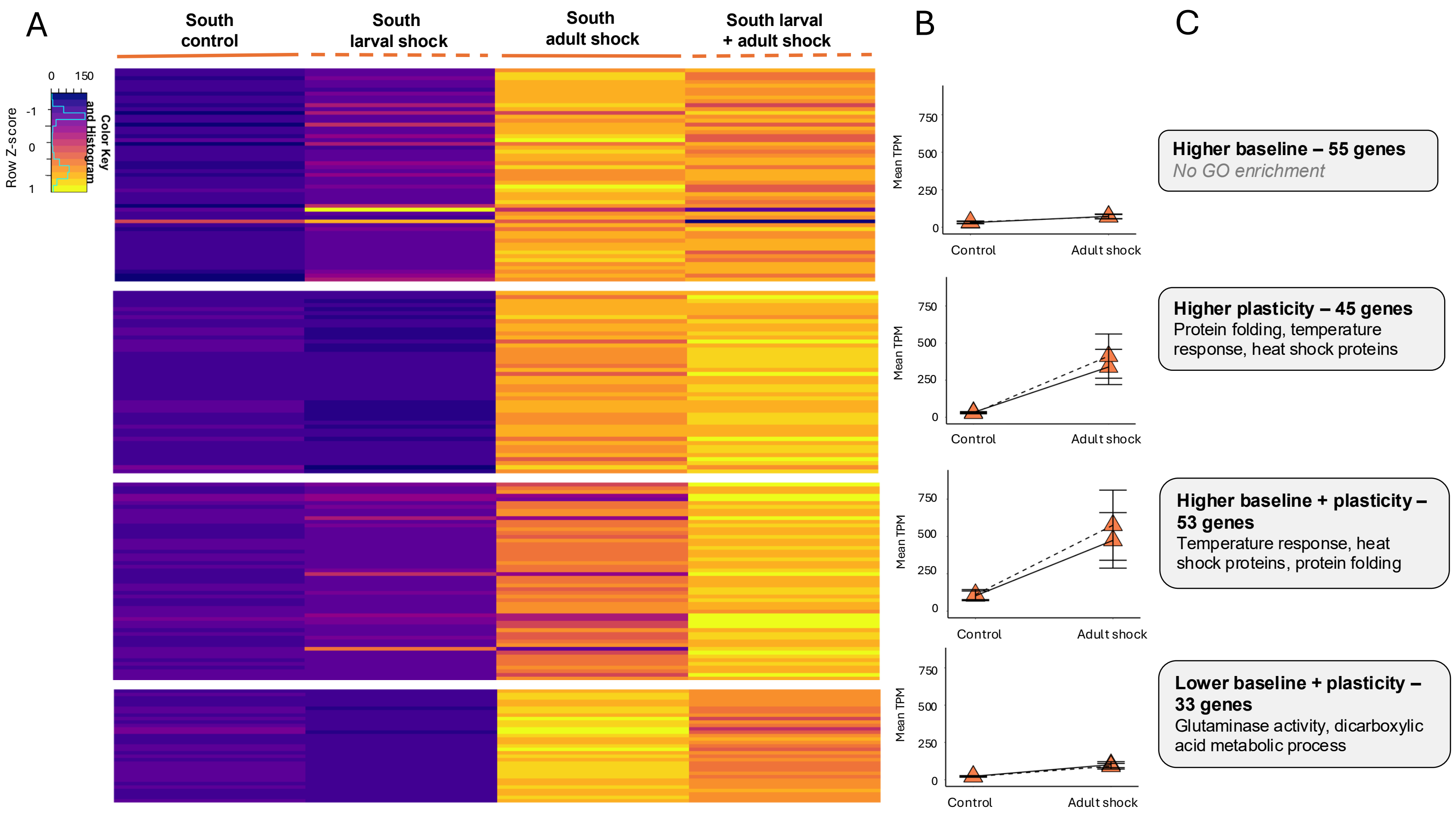


**Supplemental Figure 1** Comparison of the differentially expressed genes in the southern *Tigriopus californicus* population across four treatments (control, larval shock, adult shock, and larval + adult shock). A) Expression heat map of the 186 genes that are significantly up regulated in copepods exposed to adult heat shock as compared to controls. The mean TPM (transcripts per million) value was used for each population and treatment combination (calculated using three biological replicates) to calculate a relative expression value (Z-score). The categories were determined by 1) comparing the expression level of each gene in the larval shock treatment to the control treatment (i.e., if the larval shock treatment had a higher baseline expression) and 2) the difference in expression between control and adult shock and larval shock and larval + adult shock treatments (i.e., the difference in slope, indicating higher plasticity). B) Reaction norm plots illustrating the mean expression level (measured in TPM) within each category and across the four treatment combinations. The x-axis indicates adult treatment, and the line type indicates larval treatment (solid line indicates control, dashed line indicates larval shock). Error bars indicate standard error. C) A summary of the category, the number of genes within the category, and the gene ontology enrichment analysis indicating terms were significantly enriched within that category.
